# Light exposure induces phenotypic plasticity of the upside-down jellyfish *Cassiopea* and its endosymbiotic dinoflagellates

**DOI:** 10.1101/2024.04.08.588630

**Authors:** Rebecca Salas, Colin J Anthony, Bastian Bentlage

## Abstract

The upside-down jellyfish, *Cassiopea*, is an increasingly popular model organism gaining prominence for both its endosymbiotic dinoflagellates from the family Symbiodiniaceae and its behavioral changes of bell pulsations associated with environmental cues. Pulsation provides a unique window into the host’s response to environmental conditions, a typically difficult to access component of other symbiotic cnidarians. Pulsation has also been hypothesized to play a regulatory role on the endosymbiotic assemblage, but the magnitude of this regulatory effect is not well understood. Here, we used two light-acclimation experiments to help disentangle the complex phenotypic responses of the cnidarian host and its endosymbiotic dinoflagellates. The first experiment examined the phenotypic plasticity (size, behavior, color) of *Cassiopea* sp. in response to repeated ambient light acclimation trials to determine the rate and magnitude of phenotypic plasticity. The second experiment compared the acclimation response of jellyfish across three experimental groups to test whether a variable environment and resulting short acclimation times destabilized the host-endosymbiont relationship. Our goal was to identify covarying host-endosymbiont phenotypes to gain new insights into the dynamics of this relationship. We employed flow cytometric phenotypic profiling for high-throughput phenotypic characterization of endosymbiotic dinoflagellates in addition to pulse-amplitude modulated (PAM) fluorometry to characterize photosynthetic efficiency (F_v_/F_m_). Host phenotypes responded predictably to light-dark cycles, and stabilized after nine to twelve days of exposure to consistent light conditions. However, disruption of this acclimation period affected both the host’s circadian rhythm and the endosymbionts’ phenotypic profile. We also found evidence that phenotypic responses of the host and endosymbionts were generally decoupled, indicating a stronger regulatory response of light conditions on phenotypes than possible host-regulatory strategies on the endosymbiotic assemblage. This study provides unique insights into the acclimation strategies of upside-down jellyfish, an emerging model for the study of cnidarian-dinoflagellate symbiosis.

**Highlights:**

- *Cassiopea* behavior and color respond predictably to changing light conditions
- Inadequate acclimation time destabilizes the host’s circadian rhythm and causes unique phenotypic characteristics of the endosymbionts
- Light may be a stronger influence on host and endosymbiont phenotypes than host-endosymbiont relationships

## 1. Introduction

Upside-down jellyfish (Genus *Cassiopea*) are scyphozoan jellyfish that lie on soft-bottom substrates, typically in mangroves or other tropical and subtropical coastal waters, with their oral arms facing towards the water surface. Water-surface facing oral arms and digitate cirri facilitate photosynthesis of endosymbiotic dinoflagellates in the family Symbiodineaceae (here-in referred to as either ‘endosymbionts’ or ‘Symbiodiniaceae’), which supply nutrients to the jellyfish host, while the pulsation of their bells simultaneously pulls nutrients from the substrate and direct it through their oral arms, facilitating heterotrophy and nutrient exchange (Medina et al., 2021). In addition to stinging cells lining their oral arms and digitate cirri, *Cassiopea* also release mucous-bound clusters of stinging cells called cassiosomes into the water column to immobilize prey (Ames et al., 2020), and then use bell pulsations to direct the prey into their mouths (Hamlet et al., 2011; Santhanakrishnan et al., 2012). Bell pulsation may also regulate the jellyfish microenvironment, incorporate endosymbionts, eliminate waste, and distribute gametes (Battista et al., 2022; Freeman et al., 2016; Hamlet et al., 2011; Hoover et al., 2021; Welsh et al., 2009). These processes make *Cassiopea* spp. important ecosystem engineers by coupling benthic and pelagic nutrient cycles (Welsh et al., 2009). *Cassiopea* spp. are also effective bioindicators, as reproduction (Klein et al., 2016), behavior (Béziat and Kunzmann, 2022), morphology (Anthony et al., 2022), and physiology (Rowe et al., 2022) reflect environmental conditions. Similarly to *Cassiopea*, corals also host Symbiodiniaceae, but this relationship is threatened by increasing sea surface temperatures, as a result of accelerating global change (Brown, 1997; Hughes et al., 2017; Magel et al., 2019; Riegl and Purkis, 2015). In contrast to reef-building corals that are generally difficult to keep in captivity for experimental purposes, *Cassiopea* spp. are easily cultured and resilient, making them effective models for cnidarian symbiosis, physiology, behavior, and development (Medina et al., 2021; Ohdera et al., 2018).

Pulsating behavior of *Cassiopea* spp. provides a unique, accessible window into the host’s response to environmental cues. Circadian clocks directly translate predictable environmental signals (e.g. light levels) into behavioral or physiological responses (Reitzel et al., 2013). Both *Cassiopea* (Nath et al., 2017) and endosymbionts (Sorek et al., 2013) display a unique reversible circadian-regulated sleep-like state (quiescence) capable of responding to unpredictable changes in the environment (Nath et al., 2017). The rhythmic pulsation of the bell serves as a quantifiable measure for tracking behavioral activity during quiescence and environmental change, and may also provide some insight into the metabolic requirements of their environment (Rowe et al., 2022). In the co-evolution of host-endosymbiont interactions, symbiotic populations within the host must be controlled in order to sustain the relationship (Rivera and Davies, 2021; Xiang et al., 2020). The behavioral plasticity of *Cassiopea* in response to environmental change has been proposed to help regulate the endosymbiotic assemblage (Battista et al., 2022; Freeman et al., 2016; Welsh et al., 2009), and the endosymbionts, directly affect the host’s resilience (Kenkel and Bay, 2018; McQuagge et al., 2023; Roth, 2014).

However, light is the direct signaling stimulus known to impact photosynthetic regulation of photo-endosymbionts (Goulet et al., 2005; Sorek et al., 2013; Xiang et al., 2015; Zaheri et al., 2019). Therefore, the relationship is inherently linked and experimentally altering light conditions has the potential to disrupt both *Cassiopea* behavior and the endosymbiotic photosystem. Studying the *Cassiopea* and endosymbiont response to light stimuli may provide direct insight into covarying metabolic processes and adaptive mechanisms for environmental change in this cnidarian-Symbiodiniaceae model system.

Quantifying the acclimation dynamics of endosymbionts remains difficult, especially if attempting to resolve multiple response levels of the host and endosymbiotic assemblage (Anthony et al., 2023a; Davies et al., 2023). Cnidarians harboring photosynthetic endosymbionts change their coloration/shade depending on light conditions, indicating some level of photosynthetic regulation of the endosymbiotic assemblage. Pulse Amplitude Modulated (PAM) fluorometry remains the most widely used methodology to characterize the physiological state of the Symbiodiniaceae assemblage. PAM quantifies photosynthetic efficiency of the Symbiodiniaceae photosystems in-hospite (Berg et al., 2020; Suwa et al., 2008; Warner et al., 1996), providing an aggregate measure of photosystem performance for the Symbiodiniaceae population. However, it does not quantify variation of individual Symbiodiniaceae cells across the endosymbiont population. Flow cytometry is one method capable of quantifying phenotypic differences between individual Symbiodiniaceae cells and across entire Symbiodiniaceae assemblages by quantifying signatures of light scatter and fluorescence associated with cell size, cell shape, LHC-associated photopigments, and antioxidant-associated autofluorescent pigments (Anthony et al., 2023a; Apprill et al., 2007; Lee et al., 2012). The resolution offered by flow cytometry may reveal if changes in host color and endosymbiont photophysiology are a product of phenotypic plasticity in single Symbiodiniaceae cells or changes in cell density(Anthony et al., 2023b; Apprill et al., 2007), as well as provide insights into the covariable relationship between host and endosymbiont.

*Cassiopea* spp. are highly plastic, resilient organisms that can survive extreme physical and environmental stress across a variety of environmental conditions (Anthony et al., 2023c; Banha et al., 2020; Ostendarp et al., 2022; Tilstra et al., 2022). However, the *Cassiopea* holobiont’s phenotypic (behavior, morphology, physiology) responses to the environment, and the relationships between response strategies, are poorly defined and understood. To better understand light-mediated acclimation strategies of upside-down jellyfish, and more broadly the cnidarian-dinoflagellate relationship, we conducted two acclimation experiments. The first experiment examined the phenotypic plasticity (size, behavior, color) of *Cassiopea* sp. in response to repeated light-dark acclimation to determine the rate and magnitude of phenotypic plasticity. The second experiment compared the acclimation response of jellyfish across three experimental groups to test whether insufficient acclimation time destabilized the relationship and improve our understanding of host-endosymbiont phenotypic relationships. We hypothesized that behavior would respond predictable to light-dark cycles, and that the disruption of acclimation may affect both the host’s circadian rhythm and the Symbiodiniaceae phenotypic profile. We also hypothesized that behavior and endosymbiont phenotypes will be positively correlated, given the proposed role of pulsation in regulating the endosymbiotic photosystem.

## 2. Materials and Methods

### 2.1 Experimental design

On April 1, 2022, twenty captive-bred, presumably clonal, *Cassiopea* sp. ephyrae were acquired from Underwater World Aquarium (Tumon, Guam). *Cassiopea* sp. ephyrae were placed in separate flow-through tanks at the University of Guam Marine Laboratory and were allowed to acclimate and grow until June 6, 2022 (67 days).

To better understand the rate and magnitude of *Cassiopea* sp. phenotypic plasticity in response to periodic environmental changes, *Cassiopea* sp. medusae (50-70 mm) were alternated between two-week periods of ambient light and two-week periods of shade from June 6 to August 12, 2022 (10 weeks). Two layers of 30% UV black shade cloth was added or removed to limit available sunlight for the shaded condition, and HOBO Pendant Temperature/Light loggers (Onset, Bourne, MA) continuously monitored light (Lux) and temperature (°C) at 15 minute intervals. During the experiment, *Artemia* nauplii were continuously cultured for jellyfish feeding using a mix of 1 L seawater, 1 L freshwater, and 0.3 g of *Artemia* eggs. Jellyfish were simultaneously fed 150 mL of two-day old *Artemia* solution twice a week. Feedings took place after data collection for the day to avoid introducing feeding-related behavioral artifacts, as *Cassiopea* spp. increase their pulsation rates when presented with prey (personal observation). Pulsation rates, bell diameter, and coloration/tint of the medusae (Section 2.2) were documented three times a week, between 11:00 and 12:00, for all individuals.

To compare acclimation responses across environmental conditions, a second experiment was completed in the same tank system for an additional three weeks (August 22 - September 12, 2021) after a ten day intermediate acclimation period to ambient tank conditions. Thirteen individuals were randomly divided into three experimental groups: ‘ambient’ (n=4), ‘shade’ (n=4), and ‘variable’ (n=5). Jellyfish in the ‘ambient’ or ‘shade’ experimental groups were maintained in there condition for the entirety of the experiment. Variable individuals were rotated from ambient light for week one, shade for the second week, and to ambient light for the third week, as determined by the acclimation rates derived from experiment one. Bell diameter, pulsation rate, and color was measured three-days a week, using the same methodologies as in the previous experiment. In addition to previous metrics, pulsations and photosynthetic efficiency were measured at night (23:00), post quiescent stabilization (cf. Nath et al., 2017).

Pulsation rate was always determined prior to determining photosynthetic efficiency to avoid behavioral artifacts caused by disturbing the quiescent state of experimental medusae. At the end of the three-week experiment, tissue samples were collected from digitate cirri to determine the cell density and phenotypic profile of the Symbiodiniaceae assemblage.

### 2.2 Host phenotyping

Pulsation rate of each individual was determined by counting the number of pulsations for one minute. An analog caliper was used to measure bell diameter to the nearest millimeter. Host paling and darkening was quantified by matching the exumbrella tint to semi-quantitative color codes (D1-D6) from the CoralWatch Coral Health Chart (Siebeck et al., 2006).

### 2.3 Endosymbiont phenotyping

Maximum quantum yield (F_v_/F_m_), representative of photosynthetic efficiency, was determined using a DIVING-PAM-II fluorometer (Walz, Effeltrich, Germany) equipped with a blue LED emitting maximum light intensity at ∼475 nm. Each individual jellyfish was measured between 23:00 and 24:00 three times a week. All measurements were taken in complete darkness after isolating each individual from other individuals (to avoid light contamination). Measurements were always taken from the center of each jellyfish after placing the jellyfish on a clean piece of water-proof paper under water.

Live tissue samples were taken from digitate filaments of each jellyfish at the end of the second experiment. To quantify cell densities of endosymbiotic Symbiodiniaceae, three photos per jellyfish were taken at 100x magnification with an Olympus BX43 microscope with a DP74-CU digital camera (Olympus, Shinjuku, Tokyo, Japan). Images were scaled with a 0.1 mm reticle calibration slide. Cells within 16 mm^2^ were counted in ImageJ (Schindelin et al., 2012).

Live tissue samples were taken from digitate filaments of each jellyfish at the end of the second experiment then immediately processed for phenotypic profiling of endosymbiotic Symbiodiniaceae. Phenotypic profiles consisted of red fluorescence (LHC-associated photopigments), green fluorescence (antioxidant-related pigments), forward scatter (cell size), and side scatter (cell shape) and were quantified using the Guava easyCyte 6HT-2L (Luminex Corporation, Austin, TX) benchtop flow cytometer following the step-by-step protocol described in Anthony et al. (2023a).

### 2.4 Statistical Analyses

Experimental groups were preliminarily checked for normality using the Shapiro-Wilk test. Subsequent parametric or non-parametric comparisons of means were calculated with ANOVA or Kruskal-Wallis tests, respectively, followed by pairwise post-hoc Tukey and Dunn’s tests, respectively, depending on the outcome of the test for normality. Results for the post-hoc tests were subsequently assigned to a statistical group (e.g. A, B, C, etc.), then mapped as a tile plot to visualize statistical similarity. Groups were assigned to different groups if a pairwise comparison yielded a p-value of less than 0.05.

For the 10-week, repeated acclimation experiment, Pearson and Spearman rank-sum tests were used to identify correlations between environment (daily light and temperature) and phenotypic traits (diameter, pulsation rate, and jellyfish color), after testing for data normality with the Shapiro-Wilks test. For the 3-week comparative acclimation experiment, principal component analysis (PCA) and non-parametric Spearman-rank sum tests were used to test for experimental group effect and relationships between all phenotypic traits, respectively.

All statistics were completed with a combination of FSA v0.9.3 (Ogle et al., 2022), rcompanion v2.4.28 (Mangiafico, n.d.), and rstatix v0.7.0 (Kassambara, 2022) packages using R version 4.2.2 in the RStudio interface. Plots were generated and modified with R using a combination of the packages ggplot2 v2.4.18 (Wickham, 2016), ggpubr v0.4.0 (Kassambara, n.d.), and the open source design software inkScape v1.2.2 (inkscape.org). Code and associated datasets are publicly available on GitHub (https://github.com/AnthonyCuog/CassiopeaPlasticity).

## 3. Results

### 3.1 Environmental conditions

In both experiments, sunlight (maximum 100,000 Lux) would warm the water (30.0 - 31.5 C), without the shade cloth, while shade cloth cooled the water (28.5 - 30.5 C) and reduced light intensity (maximum 25,000 Lux), causing light intensity and temperature to covary (R^2^ = 0.884, p < 0.001).

### 3.2 Ten-week repetitive acclimation experiment

*Cassiopea* sp. grew consistently throughout the experiment, with only a minor correlation to environment changes (R^2^ = 0.103, p < 0.001) (Figure 1A). Pulsation rates increased in the ambient and decreased in the shaded environment (R^2^ = 0.301, p < 0.001), although the behavioral response to environmental change was delayed and only became apparent after 2-3 days (Figure 1B). Individuals paled in ambient environments, and darkened in the shaded environment (rho = 0.326, p < 0.001) (Fig 1C).

**Figure 1.**
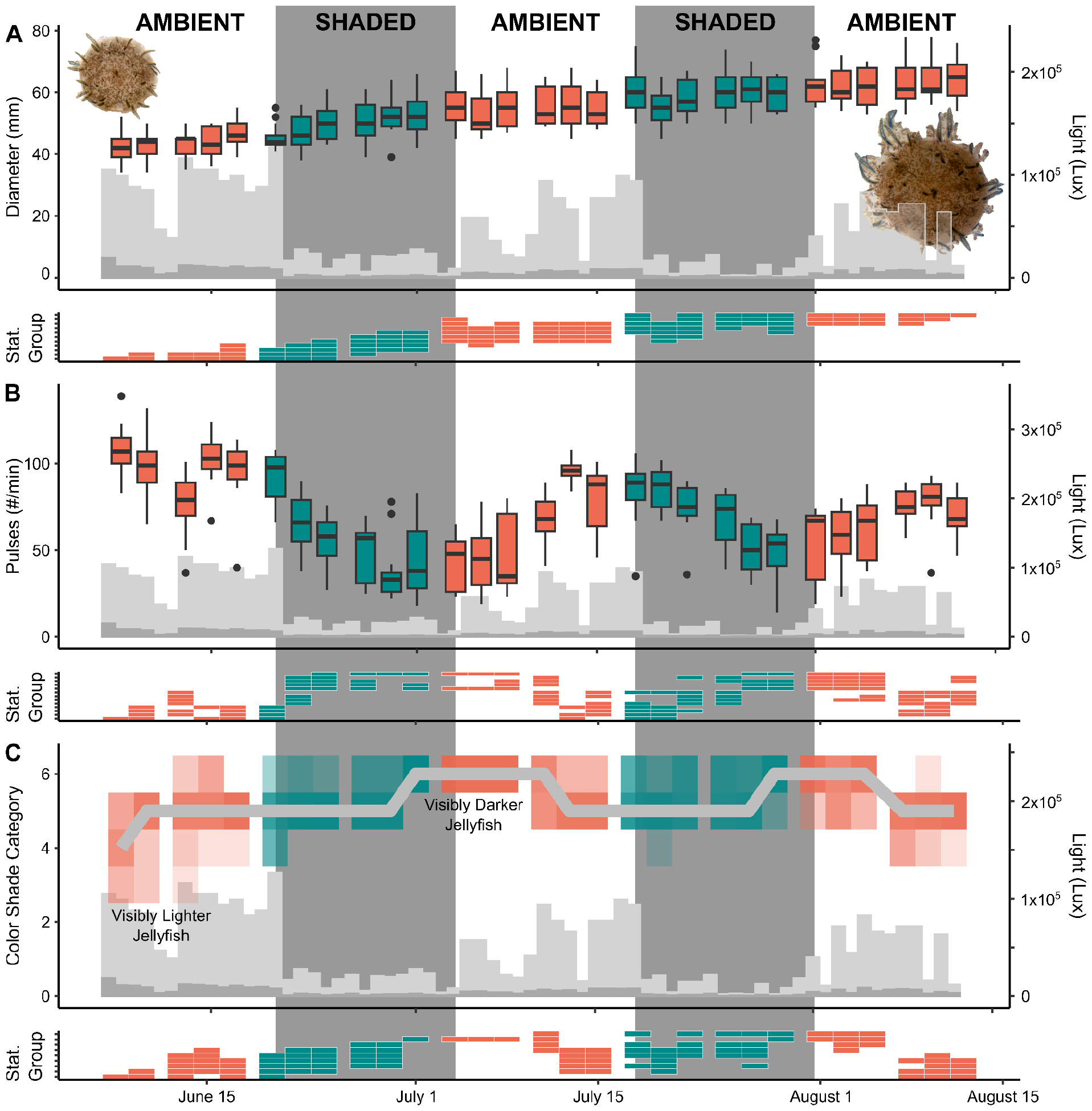
Upside-down jellyfish growth (**A**), behavior (**B**), and color (**C**) dynamics in response to 10 weeks of rotating environmental change between ambient and shaded conditions, represented by red and blue coloration, respectively. Gray plots indicate light intensity (lux). Small tile charts below each main graph indicate statistical groupings (p > 0.05) for the associated biotic response.

### 3.3 Three-week comparative acclimation experiment

Jellyfish in ambient light were at least ten millimeters larger than other experimental groups by the end of the experiment (Figure 3A). This was driven by a very slight growth of individuals in the ambient condition and a possible minor shrinkage in shaded and variably exposed jellyfish (Figure 3A).

Once individuals were placed within a stable (ambient and shaded) environmental condition, day and night pulsation rates were stable (Figure 2B). Individuals that experienced weekly environmental shifts (variable) from ambient to shaded to ambient demonstrated a destabilization of day and night pulsation rates before converging upon normal pulsation rates by the end of the final week (Figure 2B).

**Figure 2.**
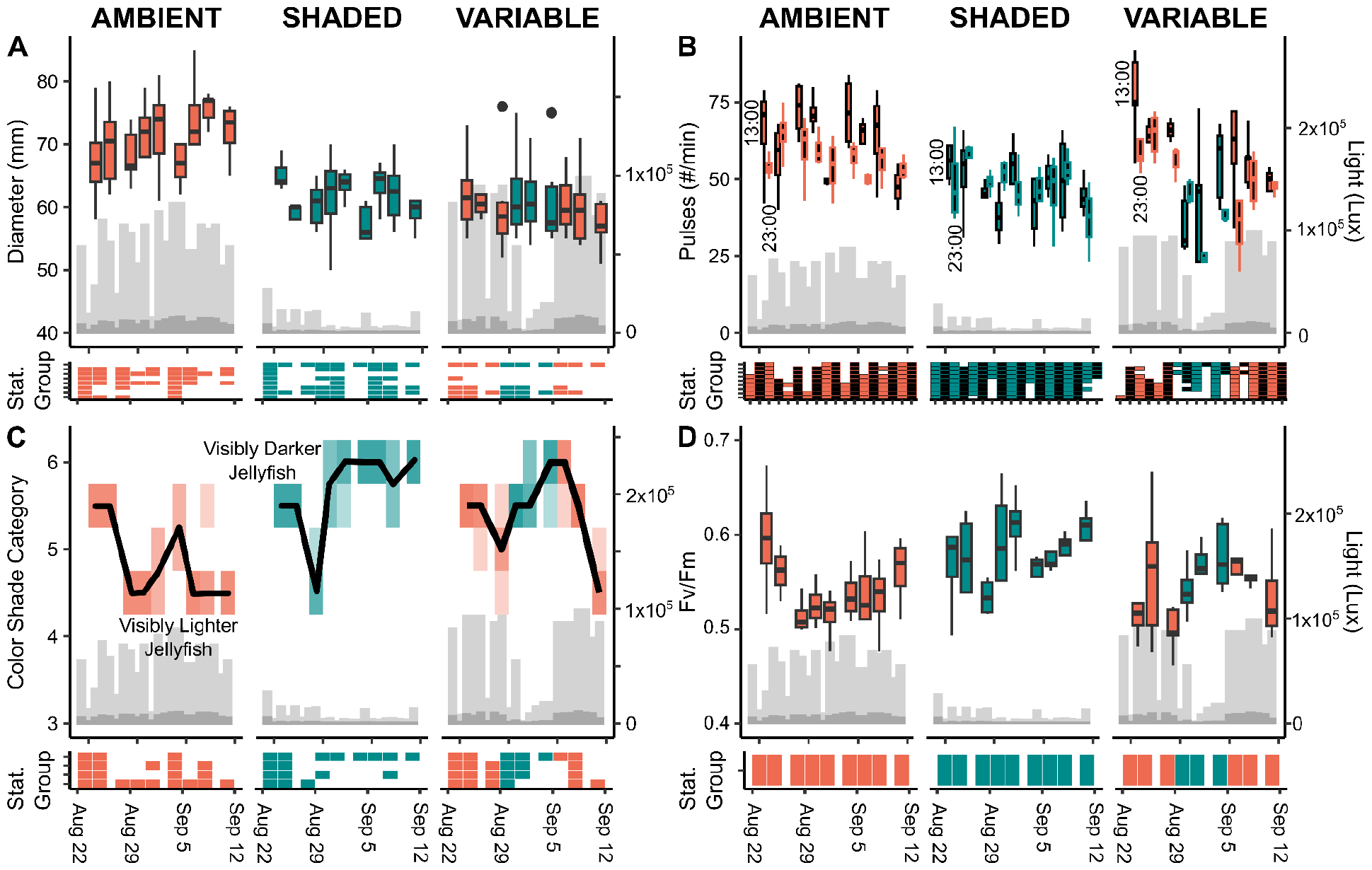
Upside-down jellyfish growth (**A**), pulsation rate in the day and night (**B**), shade (**C**), and photosynthetic efficiency (represented by Fv/Fm) (**D**) dynamics in response to 3 weeks of acclimation to 3 different experimental groups: ambient, shaded, and variable. Ambient and shaded conditions are represented by red and blue coloration, respectively. Gray plots indicate light intensity (lux). Small tile charts below each main graph indicate statistical groupings (p > 0.05) for the associated biotic response.

**Figure 3.**
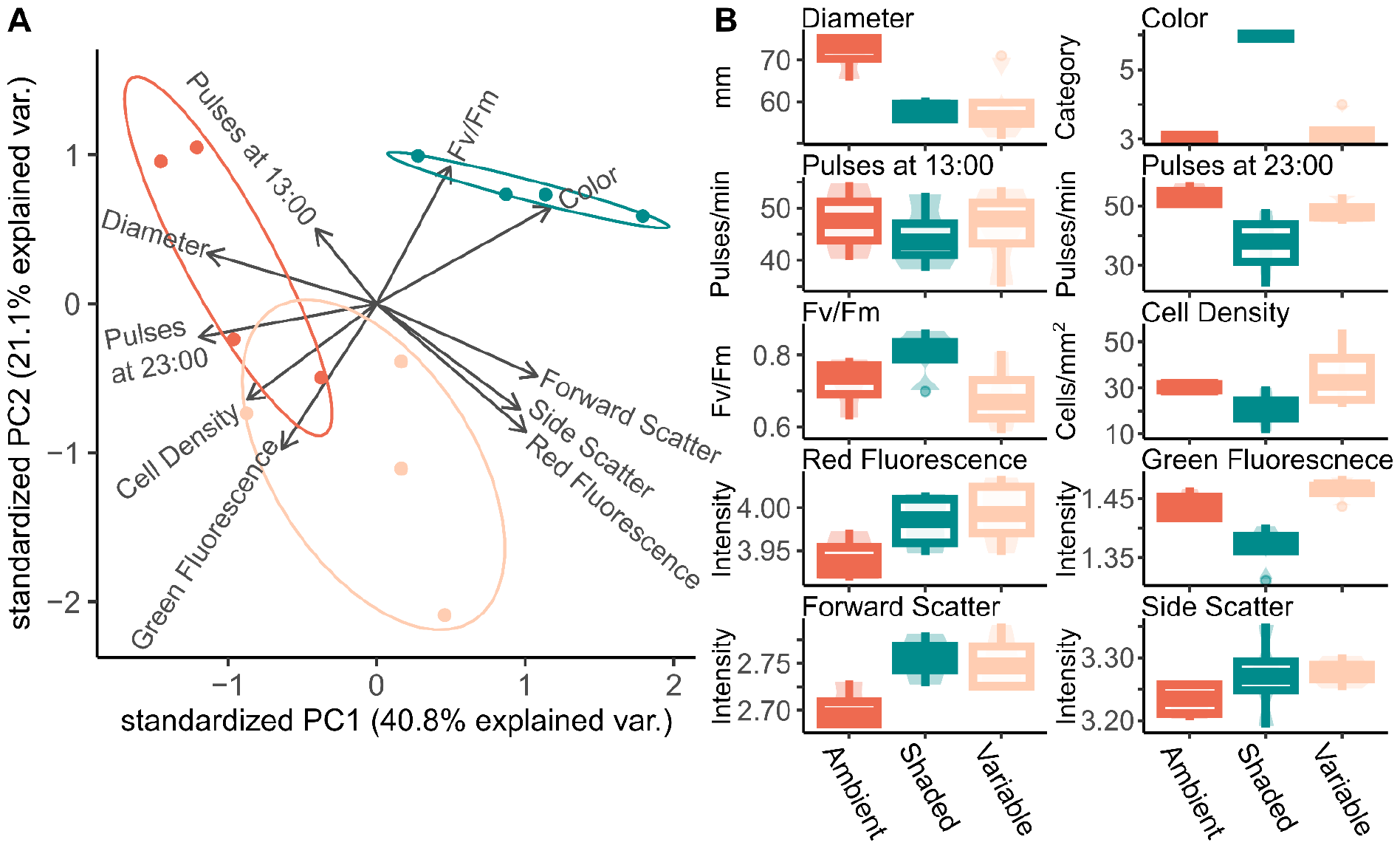
PCA of phenotypic traits (**A**) and ranges of phenotypic traits (**B**) measured at the end of the three-week comparative acclimation experiment. Colors indicate different environmental conditions (ambient, shaded, variable) jellyfish were exposed to.

Similar to the results from the 10-week repeated acclimation experiment, in the 3-week experiment, jellyfish became paler under ambient conditions and darker under shaded conditions (Figure 2C). Jellyfish from the variable environment converged upon the control groups when in both ambient and shaded conditions (Figure 2C).

Photosynthetic efficiency (F_v_/F_m_) displayed high variation within conditions, leading to no statistically significant differences in comparisons; however trends reflected the overall patterns observed in jellyfish color (rho = 0.441, p < 0.001). Generally, jellyfish in ambient conditions experienced a steep decline in photosynthetic efficiency, followed by a slow upward trend with acclimation (Figure 2D). Shaded jellyfish experienced no noticeable decline in photosynthetic efficiency (Figure 2D). Variably exposed jellyfish demonstrated an increased efficiency when shaded, followed by a decreased efficiency when under ambient conditions, although statistically significant due to high measurement variance (Figure 2D).

### 3.4 Host-symbiont phenotypic correlations

At the end of the 3-week acclimation period, phenotypic traits were measured for each individual to better understand the relationship between each metric. This included medusa diameter, jellyfish color, pulsation rate at 13:00, pulsation rate at 23:00, F_v_/F_m_ (representative of photosynthetic efficiency), Symbiodiniaceae cell density, Symbiodiniaceae red fluorescence (representative of light-harvesting photopigments), Symbiodiniaceae green fluorescence (representative of antioxidant associate photopigments), Symbiodiniaceae forward scatter (representative of cell size), and Symbiodiniaceae side scatter (representative of cell roughness) (Figure 3,4). Individuals in the shaded condition were the most distinct from ambient and variable conditions, as indicated by the PCA (Figure 3A). Jellyfish phenotypes in variable and ambient conditions still separated due to differences in Symbioodiniaceae photopigment autofluorescence, size, and roughness with phenotypic overlap caused by similar measurements for Symbiodiniaceae antioxidant-associated photopigments, cell density, photosynthetic efficiency, color, and pulsation rate (Figure 3). Shaded and variably acclimated jellyfish demonstrated a similar Symbiodiniaceae phenotypic profile with comparable photopigment, size, and roughness measurements (Figure 3).

**Figure 4.**
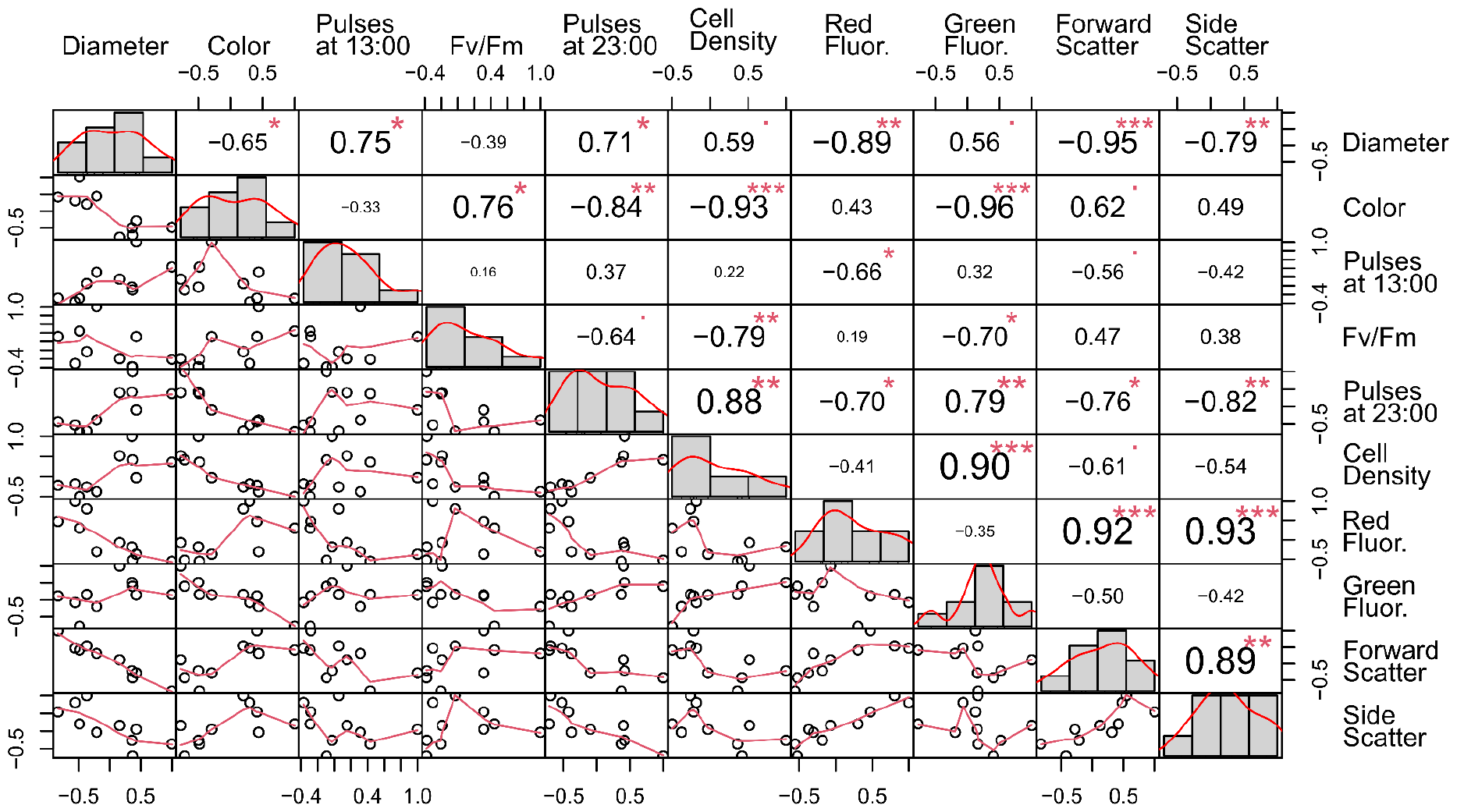
Correlations between all phenotypic traits gathered concurrently on the last day of the three-week comparative acclimation experiment. Dot plots represent the underlying data trend in relation to the two corresponding traits being compared, while numbers on the opposite side represent Spearman correlation coefficients (rho) for the compared variables. Larger numbers in size and value indicate a stronger correlation between the two variables. Red stars indicate p-values: “*” < 0.05, “**” < 0.01, “***” < 0.001. Histograms represent the underlying data structure for each phenotypic trait.

Pairwise comparisons of phenotypes revealed many strong possible relationships (Figure 4). Among the strongest (|ρ| > 0.90) were forward scatter versus jellyfish diameter, jellyfish color versus green fluorescence, color color versus cell density, green fluorescence versus cell density, red fluorescence versus forward scatter, and red fluorescence versus side scatter (Figure 4).

## 4. Discussion

Here we characterized phenotypic (behavior, morphology, physiology) changes of *Cassiopea* sp. and its associated Symbiodinaceae through cycles of repeated, short-term changes in light conditions, revealing both host and endosymbiont acclimation dynamics. The response of *Cassipea* sp. to repeated changes in light conditions (Figure 1) revealed that host behavior and coloration responded readily, with complete phenotypic stabilization occurring between days nine and twelve (Figure 1). Comparing *Cassiopea* sp. phenotypic acclimation across three, side-by-side light regimes revealed phenotypic divergence within the variable light condition, which provided insufficient acclimation time (Figure 2,3).

### 4.1 Environmental influence on host phenotype

Changes in medusa diameter was not impacted by two-week periods of decreased available light (Figure 1), but medusae grew larger in ambient light than in shaded or variable light conditions over a three-week period (Figure 2A, 3B). Previous research also indicated a higher degree of stability in medusa size during periods of environmental fluctuations (Rowe et al., 2022). Despite relative stability, medusa size is still considered a strong indicator of environmental conditions. For example, populations of smaller medusae after long periods of elevated water temperatures can indicate ‘invisible bleaching’ where jellyfish shrink, as a result of the loss of endosymbiotic Symbiodiniaceae (Morejón-Arrojo et al., 2024; Toullec et al., 2023).

In our experiment, we show that growth is affected by the light (and covarying water temperature), as evident by the significantly larger medusae observed under ambient light conditions (Figure 3B). Given the ‘invisible bleaching’ hypothesis of (Toullec et al., 2023)), if *Cassiopea* sp. individuals were heat stressed in our experiment, we would expect medusae to shrink. Given their continuous growth, jellyfish in our experiment were not heat-stressed and instead were primarily responding to changes in light conditions.

After each change in light conditions, pulsations and color took nine to twelve days before stabilizing at their new rate (Figure 1B,C). However, we find a noticeably delayed response of jellyfish phenotypes to environmental change (Figure 1B). In ambient light jellyfish pulsed faster and paled. In shaded conditions jellyfish darkened and slowed pulsation rates. This indicates a a clear correlation between light intensity and jellyfish color. These findings highlight the strong influence of light conditions on jellyfish coloration and behavior, revealing their dynamic phenotypic plasticity in response to environmental cues. This suggested a general lag period prior to full acclimation to new conditions. The lag time most likely helps predict an organism’s ability to survive in a changing climate, with a shorter lag time indicating a better ability to survive unpredictable change (Anthony et al., 2023d; Haaland et al., 2021, 2019; Stearns and Koella, 1986).

Notably, when exposed to variable conditions, *Cassiopea* sp. displayed the ability to match the pulsation and coloration of control groups in both ambient and shaded settings, showcasing their expedient acclimation to changing light conditions. However, our second experiment revealed a noticeable circadian destabilization of individuals in the ‘variable’ experimental group (Figure 2B), a metric not examined in the first experiment. Therefore, despite similar pulsation and color values during their present light condition (Figure 2), jellyfish became stressed and dysregulated without proper acclimation time of 9-12 days (c.f. Figure 1). It is possible that repeated, one-week changes in light conditions could quicken jellyfish acclimation time (Nedelec et al., 2016; Walter et al., 2013). Alternatively, environmentally induced dysregulation may cause prolonged stress that limits the jellyfish’s ability to return to physiological baseline (Berg et al., 2020). Nonetheless, we confirm our initial hypothesis and conclude that not allowing for the proper acclimation time resulted in a circadian destabilization (Figure 2B), and a distinct phenotypic profile (Figure 3A; discussed later).

### 4.2 Environmental influence on symbiont phenotype

Experiment two was also used to improve our understanding of endosymbiotic response to different light conditions. Photosynthetic efficiency (F_v_/F_m_) was highest in dark conditions, and lowest in the variable light condition. This suggests that Symbiodiniaceae exposed to variable conditions were stressed, an interpretation supported by cell density and flow cytometry-generated phenotypic profiles (Figure 3). Red and green autofluorescence signatures in *Cassiopea*-associated Symbiodiniaceae unveiled distinct responses to varying light conditions. Flow cytometry allows to characterize sample phenotypes of thousands of individual Symbiodiniaceae cells per jellyfish host individual, allowing for detection of even small but significant effects while measurements of the aggregate Symbiodiniaceae population, such as those generated by PAM fluorometry, are less sensitive (Colin Jeffrey Anthony et al., 2023a).

Shade-acclimated individuals exhibited elevated red fluorescence (indicative of light-harvesting pigments) and reduced green fluorescence (associated with antioxidants), suggesting an adjustment to the photosynthetic machinery to low light conditions. Conversely, ambient light-acclimated individuals displayed lower red fluorescence and higher green fluorescence, reflecting an acclimation response to high light conditions. For example, photosynthesis under high light conditions is well-known to increase the production of reactive oxygen species (ROS) (Foyer, 2018; Pospíšil, 2016; Rehman et al., 2016; Vetoshkina et al., 2015), which may explain the increased green fluorescence observed by us. While phenotypes of ambient, high light acclimated *Cassiopea* sp. and their Symbiodiniaceae was distinct from low light phenotypes, individuals exposed to variable light conditions displayed phenotypic traits overlapping with both ambient and shaded jellyfish. As mentioned above, green fluorescence, representative of antioxidant-associated pigments (Hennige et al., 2009; Kagatani et al., 2022; Koyama and Hashimoto, 1993; Mukherjee et al., 2013; Niedzwiedzki et al., 2022), was highest in variable light conditions (Figure 3B). We hypothesize that this elevated green fluorescence represents increases in cryptochromes or riboflavin, green fluorescing stress regulators that induce the accumulation antioxidants and central components involved in light-mediated acclimation (D’Amico-Damião and Carvalho, 2018; Deng et al., 2014; Sandoval et al., 2008; Yu et al., 2010). As the green autofluorescence signatures documented by us represent the combined signal of multiple candidate proteins and pigments, targeted experiments and characterization of these overlooked components of Symbiodiniaceae acclimation would enable more in-depth insights into the mechanisms of Symbiodiniaceae stress mitigation.

### 4.3 Host-endosymbiont phenotypic relationships

After identifying two traits that responded readily to environmental change in our first experiment (pulsation and coloration; Figure 1), we set out to understand how Symbiodiniaceae responded to differences in light regimes and if these responses corresponded with observed changes in host phenotypes. Pulsations at 13:00 and medusa bell diameter were along the same axis as Symbiodiniaceae red fluorescence, forward scatter, and side scatter (Figure 3A), but in opposite directions, indicating a negative correlation between host phenotypes and symbiont phenotypes (Figure 4). This was also the axis that separated ‘ambient’ and ‘variable’ experimental groups (Figure 3A), indicating that these phenotypic characteristics are likely linked and directly affected by an unstable, variable environment. *Cassiopea* sp. individuals that finished the experiment under ambient light conditions (‘ambient’ and ‘variable’ groups) were separated from the ‘shaded’ group along an axis characterized by medusa color, pulsation at 23:00, Symbiodiniaceae photosynthetic efficiency (F_v_/F_m_), Symbiodiniaceae cell density, and Symbiodiniaceae green fluorescence, suggesting that these traits were most affected by the light conditions experienced by individuals directly prior to sampling at the end of the experiment (‘ambient’ versus ‘shaded’ groups).

The observed behavioral changes challenge our original hypothesis that *Cassiopea* spp. pulsation regulates photosynthesis of Symbiodiniaceae. If the increased bell pulsation was regulating the photosystem (Verde and McCloskey, 1998; Welsh et al., 2009), we would most-likely expect a positive correlation between pulsation and LHC-associated fluorescence and cell shape. Instead, we see limited indication at all that pulsation rate was correlated at all with endosymbiont phenotypes (Figure 4). Additionally, medusae were largest in ambient light conditions, which was accompanied by a higher pulsation rate and higher Symbiodiniaceae cell densities (Figure 3). However, larger jellyfish have previously been shown to pulse less frequently and harbor Symbiodiniaceae at lower cell densities than smaller jellyfish (Fitt et al., 2021; Verde and McCloskey, 1998), which is not the relationship observed here (Figure 4). This confirms that our results are representative of the environmental condition’s effect on either the host or the endosymbiont, independently. Our results do not support host-endosymbiont linked phenotypic relationships. While it’s conceivable that photosystem regulation may be an auxiliary effect of pulsation, based on our research and previous research, pulsation most like primarily serves substrate attachment, and heterotrophy (Battista et al., 2022; Durieux et al., 2021; Hamlet and Miller, 2012; Santhanakrishnan et al., 2012), before photosynthetic regulation.

Contrary to the assumption that Symbiodiniaceae cell densities ought to correspond to darker coloration of the host jellyfish (Brown, 1997; Jones, 2008), cell density was lower in darker individuals (Figure 3). This indicates that jellyfish color may reflect other aspects of the Symbiodniaceae assemblage, or host proteins. For example, we did find that jellyfish color and photosynthetic efficiency (F_v_/F_m_) were correlated with changes in cell phenotypes (Figure 4), indicating a potential role of increased cell size, shape, or pigmentation rather than symbiont cell densities in influencing host coloration. Although we have evidence for the decoupled relationship between host color and Symbiodiniaceae cell density, it remains difficult to identify the source of host color. For example, all of our measurements of the Symbiodiniaceae assemblage were derived from characterizing the community within the oral filaments, but Symbiodiniaceae reside throughout *Cassiopea* sp. including within the oral arms, vesicular appendages, and exumbrella. All can contribute to host coloration. Additionally, there may be another component contributing to plasticity of host coloration, such as previously identified white granules that may play a role in photosynthetic acclimation (Lyndby et al., 2023). Focused research relating host phenotype to symbiont phenotype and localization in tissues will improve our understanding of this important relationship.

## Acknowledgments

We would primarily like to thank NSF INCLUDES SEAS Islands Alliance and Guam NSF EPSCoR for providing research funding and academic infrastructure that made this research possible. We would like to thank Jonathan “Nanny” Perez for helping us collect jellyfishes for our preliminary research, UnderWater World aquarium in Tumon, Guam for providing clonal jellyfish, MacKenzie Heagy for her mentorship and wet lab assistance, and Loreto Paulino Jr. for contributing his data handling expertise.

## CRediT authorship contribution statement

**Rebecca Salas:** Conceptualization, methodology, software, formal analysis, investigation, data curation, writing—original draft preparation, writing—review and editing, visualization. **Colin J Anthony:** Conceptualization, methodology, software, formal analysis, investigation, data curation, writing—original draft preparation, writing—review and editing, visualization, supervision, project administration. **Bastian Bentlage:** Conceptualization, methodology, investigation, writing—review and editing, project administration, funding acquisition

## Data availability

All data and original code is available on GitHub in a public repository: https://github.com/AnthonyCuog/CassiopeaPlasticity

